# Exploring the Interactome of PML nuclear subdomains during fatty acid stress using APEX2-mediated proximity labeling

**DOI:** 10.1101/2023.06.21.545954

**Authors:** Jordan Thompson, François-Michel Boisvert, Jayme Salsman, Graham Dellaire, Neale D. Ridgway

## Abstract

When exposed to excess fatty acids, specific cell types produce nuclear lipid droplets (nLDs) that associate with promyelocytic leukemia (PML) protein to form <u>L</u>ipid <u>A</u>ssociated <u>P</u>ML <u>S</u>tructures (LAPS) that are enriched in lipid biosynthetic enzymes but deficient in canonical proteins associated with PML nuclear bodies (PML NBs). To identify the PML interactome during lipid stress, we employed proximity-dependent biotin identification (BioID) in U2OS cells expressing PMLI and PMLII fused to the ascorbate peroxidase APEX2 and cultured in the absence or presence of oleate to enhance lipid droplet formation. The resulting interactome included proteins enriched under oleate-treated conditions, such mitogen activated protein kinase-activated protein kinase 2 (MK2), ESCRT proteins and the COPII vesicle transport proteins Sec23B, Sec24A and USO1. COPII proteins co-localized with both PML-NBs and LAPS but were selectively enriched in PML-NBs following oleate treatment. The nuclear localization of USO1 was uniquely dependent on PML expression. Thus, the APEX2-PML proximity interactome implicates PML domains in the nuclear function of a non-canonical network of COPII vesicle trafficking proteins.

**SUMMARY STATEMENT:** This is the first study to utilize APEX2 proximity labelling to identify the protein interactome of PML nuclear substructures and how interactions are modified under conditions of fatty acid-induced stress.

## INTRODUCTION

Promyelocytic leukemia (PML) is the main scaffold protein of PML-nuclear bodies (PML NBs); subnuclear domains that orchestrate cell survival pathways in response to apoptosis, DNA damage, viral infection and senescence (Attwood et al., 2020; Bernardi et al., 2008; Bischof et al., 2002; Dellaire et al., 2006). The PML gene encodes 6 nuclear isoforms that have a common RING-B-box coiled-coil (RBBC) motif and a variable C-terminal region. These PML isoforms homo-oligomerize via RBBC and SUMO-SUMO interaction motifs (SIM) to form the PML NB scaffold (Shen et al., 2006) that then recruits numerous client proteins (>150) via SUMO or SIM interactions, including transcription factors, chromatin remodeling factors, kinases, phosphatases and components of the SUMO-modifying machinery itself (Barroso-Gomila et al., 2021; Dellaire et al., 2003). In addition to PML NBs, six non-canonical PML substructures have been identified in the nuclei of cells exposed to stress stimuli, such as senescence, oncogenic transformation, nuclear envelope damage, excess fatty acids and accretion of misfolded proteins (reviewed in (Mcphee et al., 2022)). These novel PML-containing subdomains are structurally altered in response to stress and contain a different repertoire of client proteins than found in canonical PML NBs, which presumably ascribes new cell function(s) to these domains. One of these atypical PML substructure are the Lipid-Associated PML Structures (LAPS), which are comprised of a nuclear lipid droplet (nLD) surrounded by PML, CTP:phosphocholine cytidylyltransferase α (CCTα), Lipin1, and other lipid metabolic enzymes and proteins (Lee et al., 2020; Ohsaki et al., 2016). LAPS are formed during exposure of cells to excess fatty acids but their role in the cellular response to this external stress is not fully understood.

Eukaryotic cells utilize fatty acids as an energy source and for the synthesis of membranes but when demands are met, excess is stored as triacylglycerol (TAG) into cytosolic lipid droplets (cLD). In contrast, nLDs are less abundant than cLDs, smaller in size, and restricted to specific cell types. nLDs were first identified in mouse and human hepatocytes (Layerenza et al., 2013; Uzbekov et al., 2013), and later in Huh7 hepatoma cells (Ohsaki et al., 2016), U2OS osteosarcoma cells (Soltysik et al., 2021; Lee et al., 2020), yeast (Romanauska et al., 2021) and *C. elegans* (Mosquera et al., 2021). In Huh7 cells, nLDs are derived from lipid droplets in the lumen of the endoplasmic reticulum (ER) that are the precursors for assembly of very low-density lipoproteins. Under conditions of ER stress and fatty acid overload, ER luminal lipid droplets enter the type I nucleoplasmic reticulum and break through into the nucleoplasm at sites on the membrane demarcated by the PMLII isoform (Soltysik et al., 2019). During its egress into the nucleoplasm the nLD associates with PMLII to form LAPS. An alternate mechanism identified in U2OS cells that do not secrete lipoproteins involves assembly of nLDs and LAPS on the inner nuclear membrane (INM) by a mechanism akin to cLD assembly and budding from the ER (Soltysik et al., 2021). In addition to PML, nascent nLDs also associate with lipid biosynthetic enzymes, including diacylglycerol acyltransferases 1 and 2, Lipin1, and glycerol phosphate acyltransferases (Ohsaki et al., 2016; Soltysik et al., 2019; Soltysik et al., 2021; Lee et al., 2020)], indicating they arise by localized TAG synthesis on the INM. However, nLD biogenesis is independent of seipin, which is required for cLD assembly on the ER (Soltysik et al., 2021). Results with PML knockout U2OS cells showed that LAPS account for 60% of total nLDs and are depleted of canonical PML NB associated proteins, such as SUMO, DAXX and SP100 (Lee et al., 2020). These results suggest that an initial step in LAPS formation involves deSUMOylating and disruption of PML NBs, and the subsequent recruitment of PML to the surface of nLDs.

The reorganization of nuclear PML protein and its association with nLDs to form LAPS is likely associated with large scale changes in the interactome of PML subnuclear domains that could attenuate or generate a new signalling hub in response to fatty acid-induced lipid stress. Thus, to better understand the functional and dynamic relationships between PML NBs and LAPS we sought to characterize the changing interactome of PML during lipid stress. To specifically address this gap in knowledge, we employed APEX2-mediated proximity-dependent biotin identification (BioID) to characterize the interactome of PMLI and II in the context of naive and oleate-treated U2OS cells. Quantitative mass spectrometry of streptavidin-affinity purified protein revealed 64 PMLI- and PMLII-interacting proteins, many of which were enriched under oleate-treated conditions. Sec23B, Sec24A and USO1 interacted with PML NBs and LAPS, highlighting a non-canonical nuclear function for these well-known COPII coat and adaptor proteins in the ER-Golgi secretory pathway.

## RESULTS

### APEX2-PMLI and APEX2-PMLII localize to PML NBs and LAPS

The interactome of PML NBs and LAPS was investigated by biotin proximity labeling of these structures in U2OS cells using APEX2 fused to the N-terminus of PMLI and PMLII (referred to hereafter as APEX2-PMLI and APEX2-PMLII, Fig. 1A). APEX2 with an NLS fused to its C-terminus was used to control for non-specific nuclear interactions (referred to hereafter as APEX2-NLS). All three constructs have an internal HA tag to visualize expression by confocal microscopy and immunoblotting in transiently transfected U2OS cells, which were chosen for the study because they express abundant PML NBs and LAPS (Lee et al., 2020; Ohsaki et al., 2016). APEX2-PMLI, and APEX2-PML-II and APEX2-NLS were expressed in U2OS cells at the expected molecular mass when probed with an HA antibody (Fig. 1B). Confocal immunofluorescence was used to determine if APEX2-PMLI and -PMLII associated with PML NBs and LAPS (Fig. 1C and D). In transiently transfected U2OS cells both APEX2-PMLI and APEX2-PMLII were localized to nuclear puncta that were positive for HA and PML (Fig. 1C and D). Treatment of these cells with oleate for 24 h resulted in the appearance of BODIPY493/503-positive nLDs that had HA-tagged fusion proteins on their surface (Fig. 1C and D) indicating that both APEX2 probes were incorporated into PML structures.

**Figure 1.**
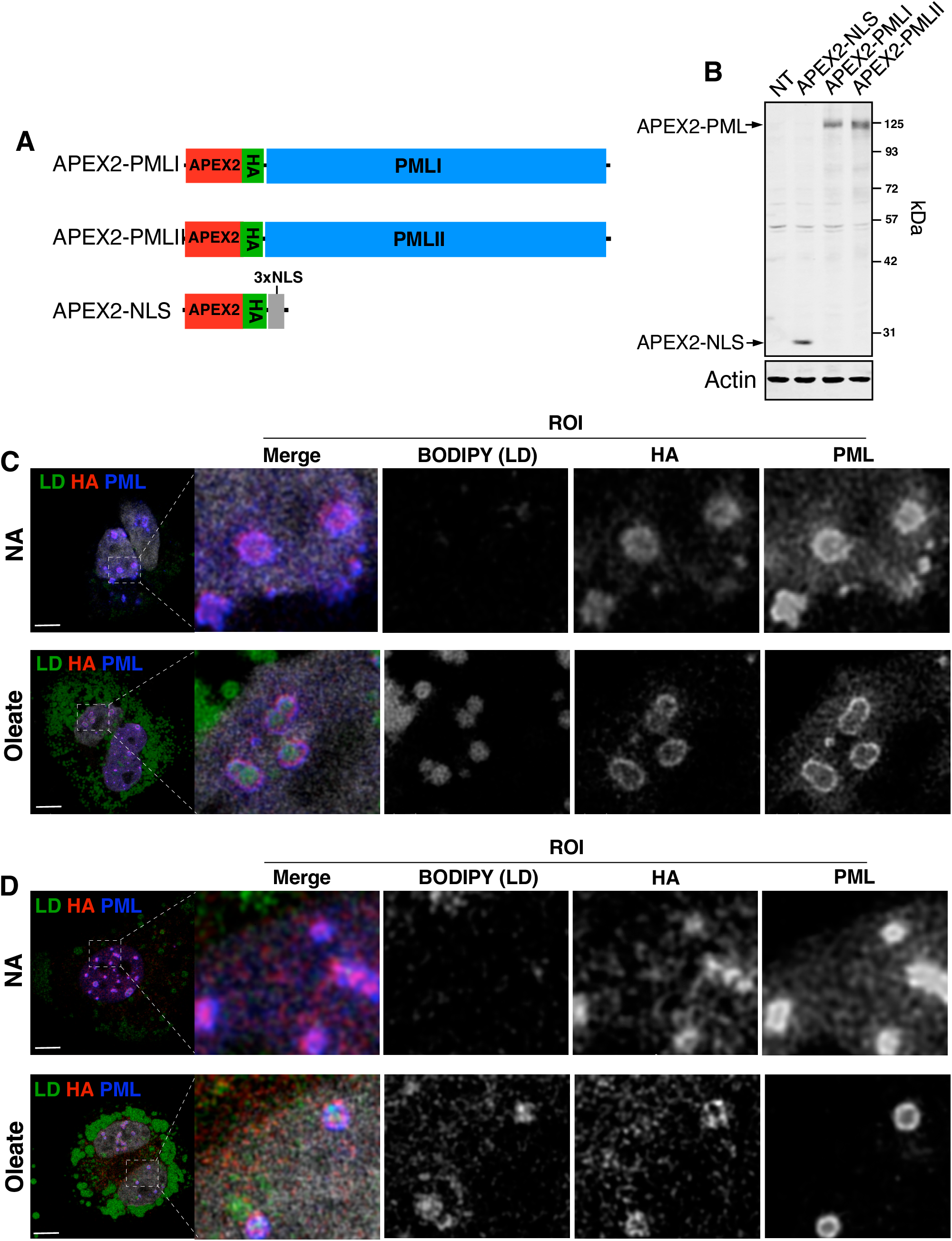
APEX2-PMLI and -PMLII localize to LAPS. A, structures of APEX2-PML and APEX2-NLS fusion proteins. B, lysates of U2OS cells transiently expressing APEX2-PMLI, APEX2-PMLII or APEX2-NLS were immunoblotted with a HA antibody (NT, non-transfected). C and D, U2OS cells transiently expressing APEX2-PMLI (panel C) or -PMLII (panel D) were cultured in media with no addition (NA) or 500 μM oleate for 24 h. Cells were immunostained with antibodies against HA and PML, followed by AlexaFluor-594 and −647, respectively. LDs were visualized with BODIPY493/503 and nuclei were stained with Hoechst (grey). In the Region of Interest (ROI), the channels were split and converted to greyscale (bar, 5 μm).

Next, we determined whether APEX2-PML proteins facilitated the biotin labeling of PML NBs and LAPS. U2OS cells treated with oleate and transiently expressing APEX2-PMLI or -PMLII were treated with biotin-phenol followed by H2O2, and biotinylated proteins were visualized by immunofluorescence with streptavidin-AlexaFluor-488 (Fig. 2A). Fluorescence was restricted primarily to the nuclei where it was concentrated on LipoTox Red-positive nLDs. There was also biotin labelling of punctate structures that lacked LipoTox Red corresponding to PML NBs. These puncta were also evident in cells not treated with oleate (results not shown). The APEX2-NLS control protein was diffusely localized to the nucleoplasm but not to the surface of LAPS or PML NBs (Fig. 2B). SDS-PAGE revealed that U2OS cells expressing the three APEX2-fusion proteins and cultured in the absence or presence of oleate were positive for biotinylated proteins detected with a streptavidin-conjugated fluor (Fig. 2C).

**Figure 2.**
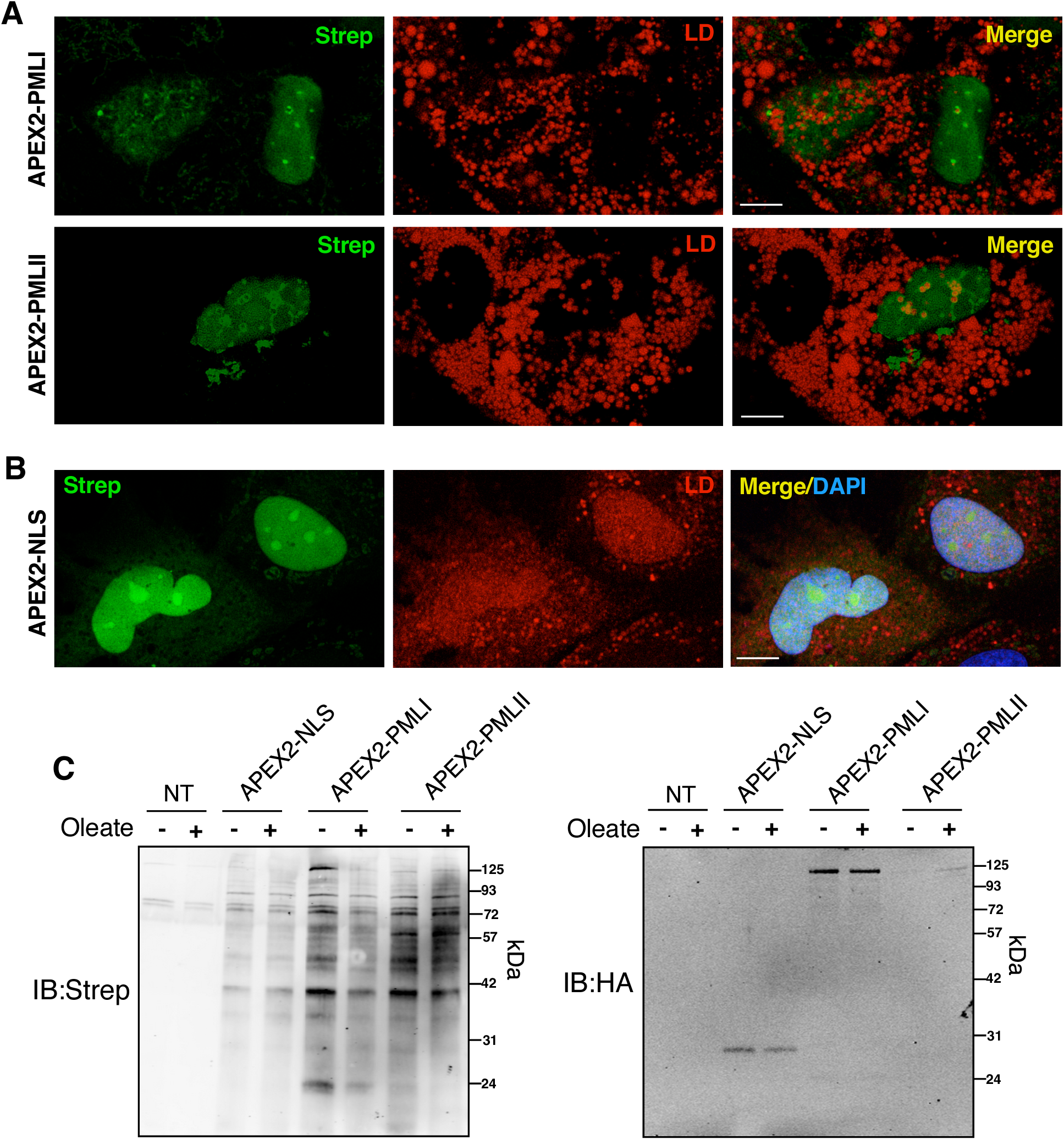
Biotinylation of LAPS proteins by APEX2-PMLI and -PMLII. A, U2OS cells expressing APEX2-PML1 or -PMLII were incubated with 500 μM oleate for 24 h followed by biotin-phenol (500 μM) and H2O2 (1 mM). Biotinylated proteins were visualized with stretavidin-AlexaFluor-488 (Strep) and LDs with LipidTOX neutral red (bar, 5 μm). B, U2OS cells expressing APEX2-NLS were treated and immunostained as described in panel A (bar, 5 μm). C, U2OS cells expressing the indicted proteins were treated with or without 500 μM oleate for 24 h and biotinylated as described in panel A. Total cell lysates were resolved by SDS-PAGE and immunoblotted with streptavidin-conjugated LiCor680IR (Strep) and an HA monoclonal antibody.

### The APEX2-PML interactome of PML NBs and LAPS

Since the APEX2-PML proteins were localized to nuclear bodies and LAPS similarly to their endogenous counter parts, BioID was used to establish the interactome of these structures. U2OS cells expressing APEX2-PML-I, APEX2-PML-II, or APEX2-NLS were treated with or without 500 μM oleate for 24 h, followed by biotin-phenol and H2O2. Prior to quantitative mass spectrometry analysis of biotinylated proteins, the expression of fusion proteins and efficiency of the biotinylation reaction was confirmed in cell lysates by immunoblotting with a HA antibody and a streptavidin probe (see Fig. 2C). High confidence APEX2-PMLI and -PMLII interactors were identified using SAINT score analysis and filtered for non-specific interactions against the APEX2-NLS and CRAPOME data sets. A total of 64 proteins interacted with both PML isoforms in PML NBs (no addition) and LAPS (oleate) and were sorted into a dot plot to illustrate the isoform- and condition-specific interactions (Fig. 3A). Although many interacting proteins were shared between the two PML isoforms, more high confidence interactions were identified for APEX2-PMLI. Approximately 75% of the interactors are nominally cytoplasmic or localized to cytoplasmic organelles, with the remainder designated as nuclear-localized (Fig. 3B). Well-known PML NB and LAPS interactors identified by other methodologies were not represented in our set suggesting that biotin proximity labelling targets a unique subset of interactions in these PML structures. To facilitate identification of common networks, proteins that interacted with both PMLI and PMLII in the presence (Fig. 3B) and absence of oleate (Fig. 3C) were pooled and analysed using STRING (disconnected proteins were excluded from the network based on confidence using all active interaction sources). The resulting networks highlighted 2 interconnected clusters: 1) Endosomal sorting complexes required for transport (ESCRT) proteins such as TSG101, STAM and STAM2 and 2) COPII vesicular coat proteins and tethers such as Sec23B, Sec24A, Sec24B, USO1 and Sec23IP. These interaction networks were evident with both PMLI and PMLII and under control and oleate-treated conditions but were more robust for APEX2-PMLI in oleate-treated cells.

**Figure 3.**
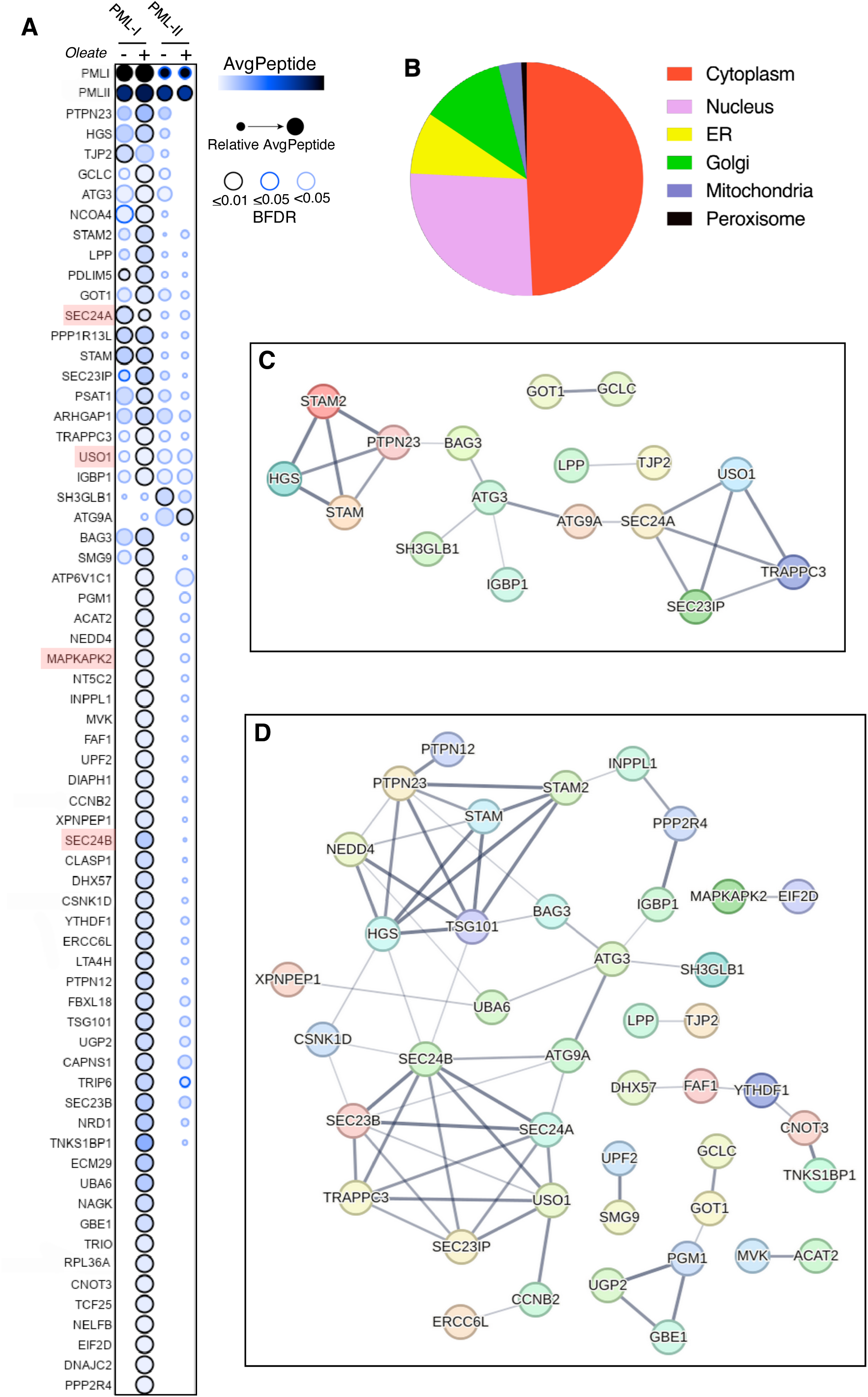
PMLI and PMLII interaction network in control and oleate-treated U2OS cells. A, dot matrix showing the average peptide counts, relative average peptides, and Bayesian false discovery rate (BFDR) for APEX2-PMLI and -PMLII interactors. Proteins selected for further characterization are highlighted in red. B, predicted cellular localization of the interactome shown in Panel A. C and D, String protein interaction networks for PMLI plus PMLII under control (panel C) and oleate-treated (panel D) conditions. Shown are proteins with validated interactions based on medium confidence setting (disconnected nodes were excluded).

### Effect of PML knockout and oleate treatment on p38-MK2 signalling

Numerous proteins that interacted with both PMLI and PMLII were specifically enriched in oleate-treated cells, potentially pointing to LAPS-specific interactions. One of these was mitogen activated protein kinase-activated protein kinase 2 (MAPKAPK2, referred to hereafter as MK2), which is activated by p38 and phosphorylates heat shock protein 27 (HSP27) and receptor-interacting serine/threonine-protein kinase 1 (RIPK1) (Jaco et al., 2017; Wu et al., 2007). MK2 also stabilizes tumor necrosis factor mRNA by phosphorylating proteins that bind the 3’AU-rich elements (Neininger et al., 2002). The potential involvement of the p38-MK2 pathway in LAPS function is suggested by the recent finding that PML interacts with p38 and MK2, leading to inhibition of MK2 and RIPKI phosphorylation and the activation of necroptosis and apoptosis (Chen et al., 2021).

We were unable to identify the MK2 isoform that interacted with PMLI and -II in the APEX2 screen but chose full-length MK2-II isoform (400 amino acid) for further analysis since it contains a C-terminal NLS while the truncated MK2-I isoform (370 amino acid) does not (Zu et al., 1994). To validate the MK2 interaction with PML, U2OS cells transiently expressing MK2-II-FLAG were imaged by confocal microscopy after oleate treatment for 24 h (Fig. 4A). MK2-II-FLAG was diffusely expressed throughout U2OS cells but appeared to concentrate around cLDs and LAPS that contained endogenous PML (Fig. 4A). Further attempts to confirm that PML and MK2 physically interacted using co-immunoprecipitation and proximity ligation assays (PLA) were not successful due to poor antibody specificity. As an alternate approach we investigated whether activation of the p38-MK2-HSP27 signalling pathway with anisomycin was affected by PML knockout or oleate treatment. U2OS cells have a functional p38-MK2 pathway based on increased phosphorylation of MK2-T334 and HSP27-S82 after treatment with the p38 activator anisomycin (Fig. 4B). Furthermore, the p38 inhibitor SB-203580 (SB) blocked phosphorylation of both MK2-T334 and HSP27-S82, while the MK2 inhibitor PF-3644022 (PF) blocked only HSP27-S82 phosphorylation. Next, we determined whether anisomycin activation of the p38-MK2-HSP27 signaling cascade was affected in U2OS PML KO cells or by oleate treatment (Fig. 4C). Quantification of immunoblots (for example Fig. 4C) revealed that anisomycin increased p38-pT180-pY182 approximately 5-fold in U2OS cells regardless of oleate treatment (Fig. 4D). However, p38-pT180-pY182 was reduced slightly but significantly in PML KO cells treated with oleate. MK2-pT334 was increased 3-4-fold by anisomycin in U2OS and PML KO cells, which was unaffected by oleate-treatment (Fig. 4E). The relative effect of oleate on phospho-regulation of p38, MK2 and HSP27 in U2OS and PML KO cells is summarized in Fig. 4F; oleate treatment caused a minor reduction in p38 phosphorylation only in PML KO cells that did not translate into significant changes in activation of its downstream targets MK2 and HSP27.

**Figure 4.**
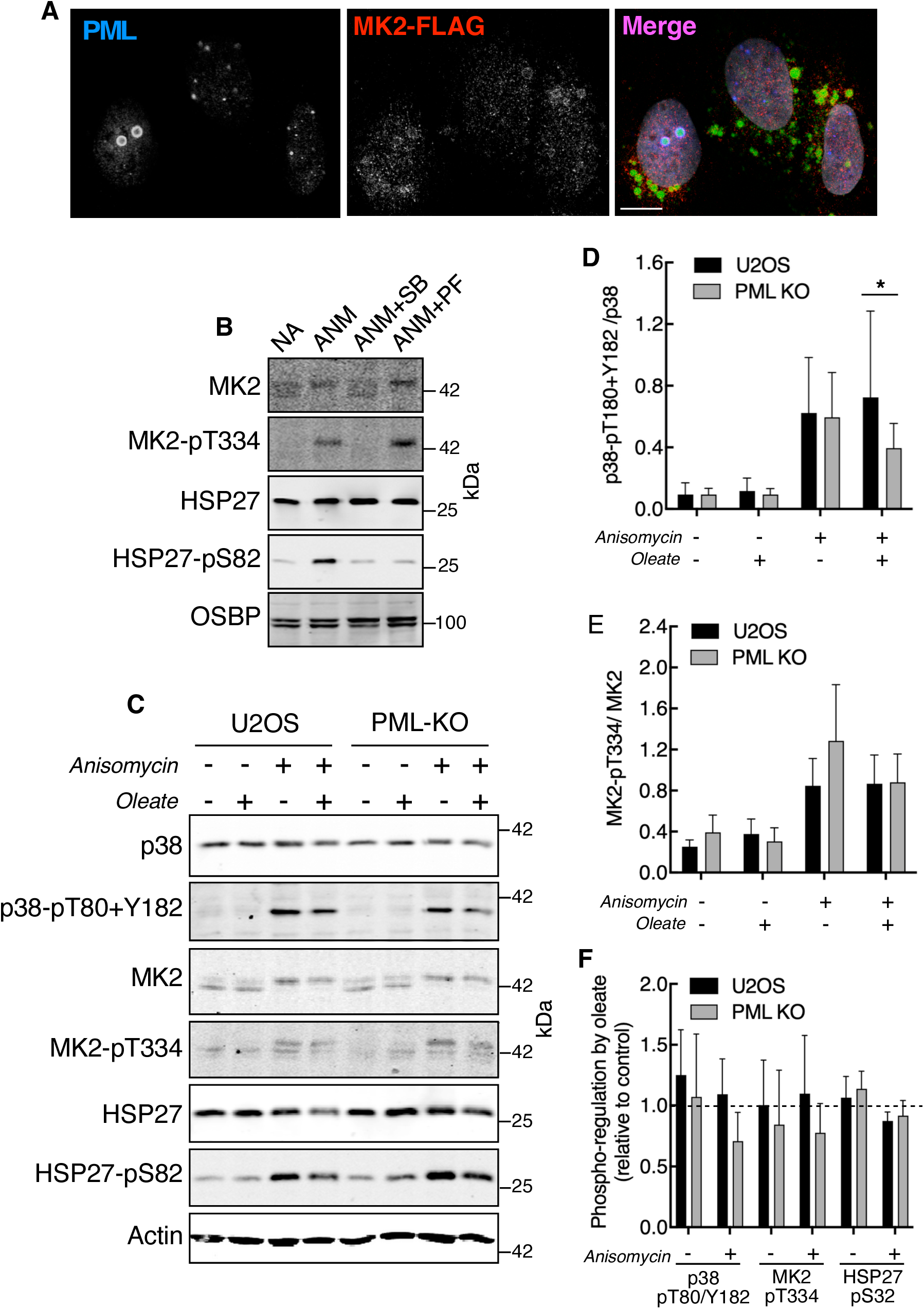
**Effect of oleate and PML knockout on the p38-MK2 signaling pathway in U2OS cells**. A, U2OS cells transiently expressing MK2-Flag (isoform 2) were treated with 500 μM oleate for 24 h and immunostained with FLAG and PML primary antibodies followed by AlexaFluor-594 and −649 secondary antibodies, respectively. LDs were visualized with BODIPY493/503 (bar, 5 μm). B, U2OS cells were pretreated for 20 mins with 10 μM SB-203580 or PF-3644022 followed by 500 nM anisomycin (ANM) for 30 mins. Cell lysates cells were resolved by SDS-PAGE and immunoblotted with antibodies against MK2, MK2-pT334, HSP27, HSP27-pS82 and OSBP (protein load control). C, U2OS and PML KO cells were cultured in media with 500 μM oleate for 24 h before treatment with 500 nM anisomycin for 30 min. Cell lysates cells were resolved by SDS-PAGE and immunoblotted with the antibodies described in panel B as well as antibodies against p38 and p38-pT180/Y182. Actin was used to normalize protein loading. D, quantitation of p38-pT180/Y182 from the immunoblots shown in panel C. Results are the means and SD of 3-6 separate experiments (*p<0.05). E, quantitation of MK2-pT334 from the immunoblots in panel C. Results are the means and SD of 3-6 separate experiments. F, the effect of oleate treatment on phosphorylation of p38, MK2 and HSP27 in U2OS and PML KO cells in response to anisomycin treatment was quantified from the immunoblots in panel B. Results are expressed relative to cells that were cultured without oleate and are the means and SD of 3-6 separate experiments.

### Interaction of COPII transport proteins with PML NBs and LAPS

A intriguing feature of the PML interactome was the association of ESCRT and COPII transport proteins with PMLI and II (Fig. 3). We chose to validate and functionally analyse the COPII network by assessing the localization of Sec23B, Sec24A and USO1 with PML NBs and LAPS using a combination of confocal immunofluorescence and PLA. Consistent with their role in COPII vesicle transport between the ER and Golgi apparatus (Aridor et al., 2001), Sec23B and Sec24A puncta were primarily localized to the perinuclear compartment of control and oleate-treated cells (Fig. 5A and B). In addition, Sec23B- and Sec24A-positive puncta were evident in the nuclei of cells cultured with or without oleate. Magnified ROIs showed potential co-localization of Sec23B and Sec24A with PML NBs and LAPS as well as nLDs devoid of PML. However, neither protein formed discrete rings or clusters on LAPS or nLDs.

**Figure 5.**
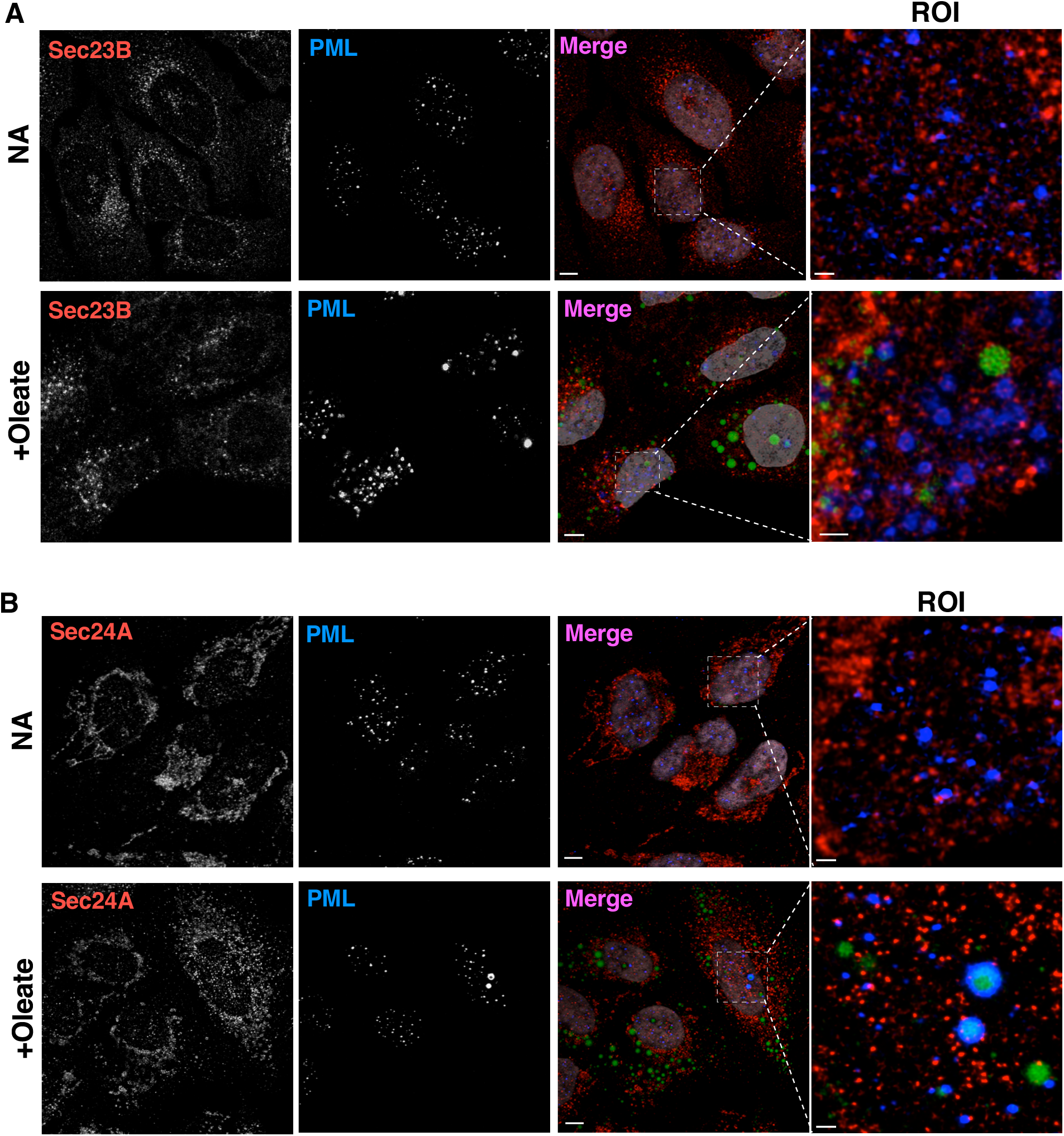
Sec23B and Sec24A-positive nuclear puncta localize to PML NBs and LAPS. A, U2OS cells were cultured in the absence or presence of 500 μM oleate for 24 h. Cells were subsequently immunostained with Sec23B and PML primary and AlexaFluor-594 and −647 secondary antibodies, respectively. LDs were visualized with BODIPY493/503 and the nuclei with Hoechst (grey) (bar, 5 μm). The region of interest (ROI) has the Hoechst signal removed (bar, 1 μm). B, U2OS cells were treated and immunostained as described in panel A but with the substitution of a Sec24A primary antibody.

PLA was used to confirm the interaction of Sec23B and Sec24A with PML. This technique involves primary antibody pairs to suspected interacting proteins, secondary antibodies with DNA tags that ligate when targets are within 40 nm, and a rolling circle DNA amplification step to enhance sensitivity (Fredriksson et al., 2002). First, we tested whether this method could be used to detect protein interactions on PML NBs and LAPS. A polyclonal antibody against DAXX, a canonical PML NB-associated protein that is depleted from LAPS (Lee et al., 2020), when combined with a PML monoclonal antibody produced numerous PLA puncta in U2OS cells (Fig. S1A) that were severely reduced by oleate treatment for 24 h (Fig. S1B and C). Secondly, a polyclonal antibody against CCTα, which associates with nLDs and LAPS in oleate treated U2OS cells (Lee et al., 2020), coupled with a PML antibody produced numerous PLA puncta in untreated U2OS cells (Fig. 6A), indicating that CCTα also associates with PML NBs. In oleate-treated U2OS cells, there were strong PLA signals on LAPS but an overall reduction in puncta due to coalescence of the PML and CCTα onto LAPS (Fig. 6B and C). The absence of CCTα-PML puncta in PML KO cells confirmed the specificity of the reaction (Fig. 6C).

**Figure 6.**
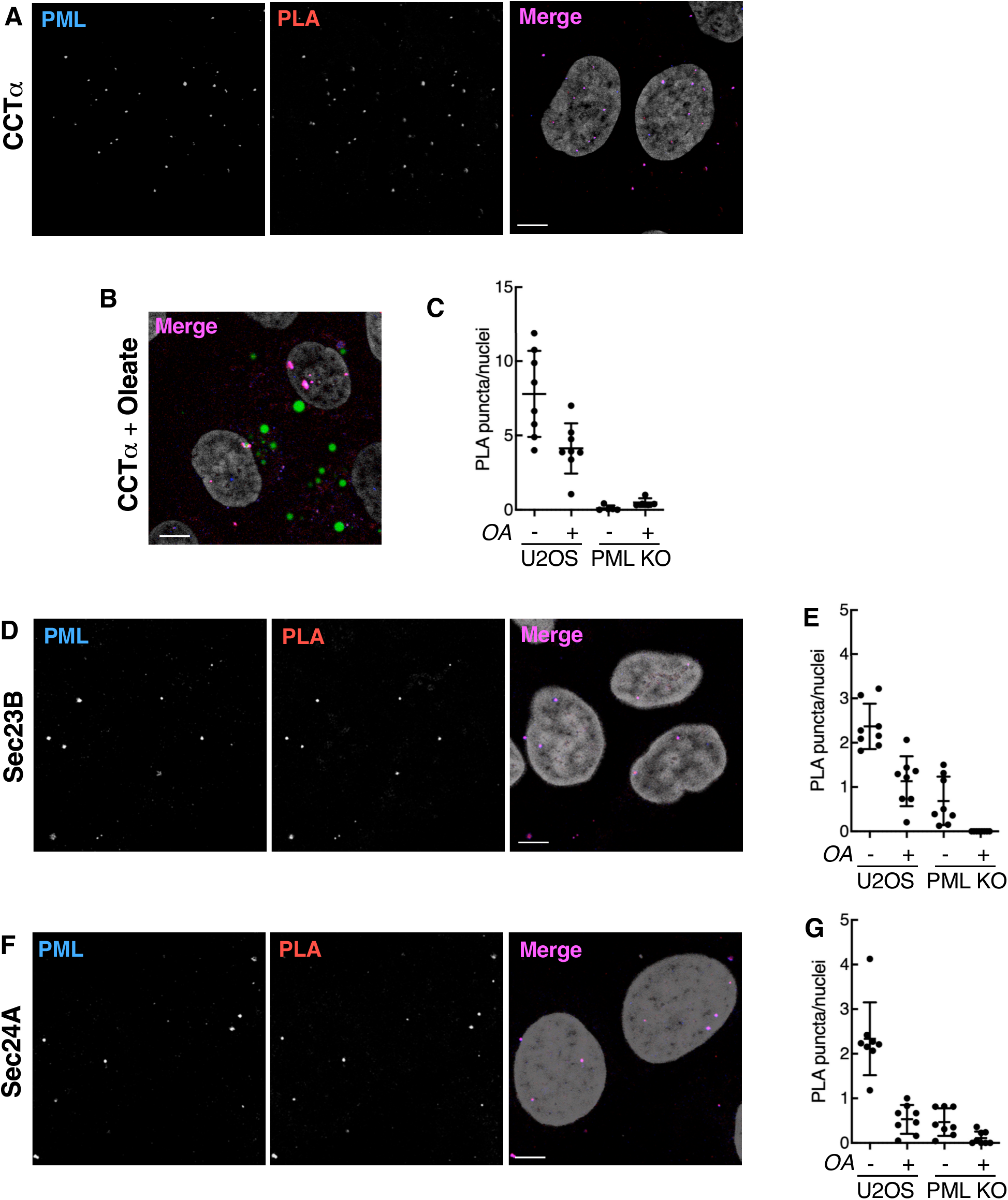
Detection of Sec23B and Sec24A interaction with PML using proximity ligation assays. A, U2OS cells were incubated with PML mouse monoclonal and CCTα rabbit polyclonal primary antibodies and subject to PLA. PML was detected with a AlexaFluor-647 secondary antibody. B, U2OS cells treated with 500 μM oleate for 24 h were incubated with CCTα and PML primary antibodies and subjected to PLA. LDs were detected with BODIPY493/503. C, PLA nuclear puncta for CCTα and PML were quantified in U2OS and PML KO cells treated with or without 500 μM oleate (OA) for 24 h. Results are the mean and SD from a representative experiment (8 fields of cell each). D, U2OS cells were incubated with PML monoclonal and Sec23B polyclonal antibodies and subjected to PLA. PML was detected with a AlexaFluor-647 secondary antibody. E, Quantification of PLA nuclear puncta for Sec23B and PML in U2OS and PML KO cells incubated in the presence or absence of oleate for 24 h. Results are the means and SD of 2 experiments (8 fields of cells each). F, U2OS cells were incubated with PML monoclonal and Sec24A polyclonal antibodies and subjected to PLA. PML was detected with a AlexaFluor647 secondary antibody. G, Quantification of PLA nuclear puncta for Sec24A and PML in U2OS and PML KO cells incubated in the presence or absence of oleate for 24 h. Results are the means and SD of 2 experiments (8 fields of cells each). Scale bar in all panels is 5 μm.

Sec23B and PML antibodies produced discrete PLA nuclear puncta in U2OS cells that overlapped with PML immunostaining (Fig. 6D). Quantification of puncta showed that oleate treatment diminished the Sec23B-PML interaction, which was partially or completely lost in PML KO cells (Fig. 6E). Similarly, PLA puncta using Sec24A and PML antibodies in U2OS cells were reduced 4-fold by oleate treatment and almost completely absent in PML KO cells (Fig. 6F and G). The number of PLA interactions detected with Sec23B, Sec24A and PML antibodies (2-2.5 per cell) was lower that with DAXX or CCTα antibodies (8-15 per cell) indicating fewer Sec23B and Sec24A interactions with PML or that methanol fixation partially disrupted PML NBs and LAPS prior to the PLA.

USO1 is a coiled-coil protein that has a canonical role in the tethering of COPII vesicle in the ER-Golgi secretory pathway (Nakamura et al., 1997; Sapperstein et al., 1995; Yamakawa et al., 1996). Immunofluorescence confocal microscopy of control and oleate-treated U2OS cells revealed the expected localization of USO1 to Golgi-like structures in the cytoplasm (Nakamura et al., 1997) as well as USO1-postitive structures that partially localized to PML NBs, LAPS and nLDs (Fig. 7A). Confocal sections through the nuclei of U2OS Clover-PML cells showed the presence of small USO1 structures on the periphery and interior of large LAPS (Fig. 7B). 3D reconstructions of a LAPS in Fig. 7B confirmed that USO1 structures were on both the exterior and interior of the LAPS apparently interacting with PML and the lipid core. The proximity of USO1 to PML structures was confirmed by PLA using a USO1 and PML antibody pair that produced numerous puncta in U2OS cells that co-localized with PML immunostaining (Fig. 7C). Quantitation of images like that shown in Fig. 7C indicated that oleate reduced PLA puncta by 2-fold, and that the signal was greatly attenuated in PML KO cells (Fig. 7D). These data point to an additional non-canonical role for Sec23B, Sec24A and USO1 in the nucleus where they interact with PML NBs and LAPS.

**Figure 7.**
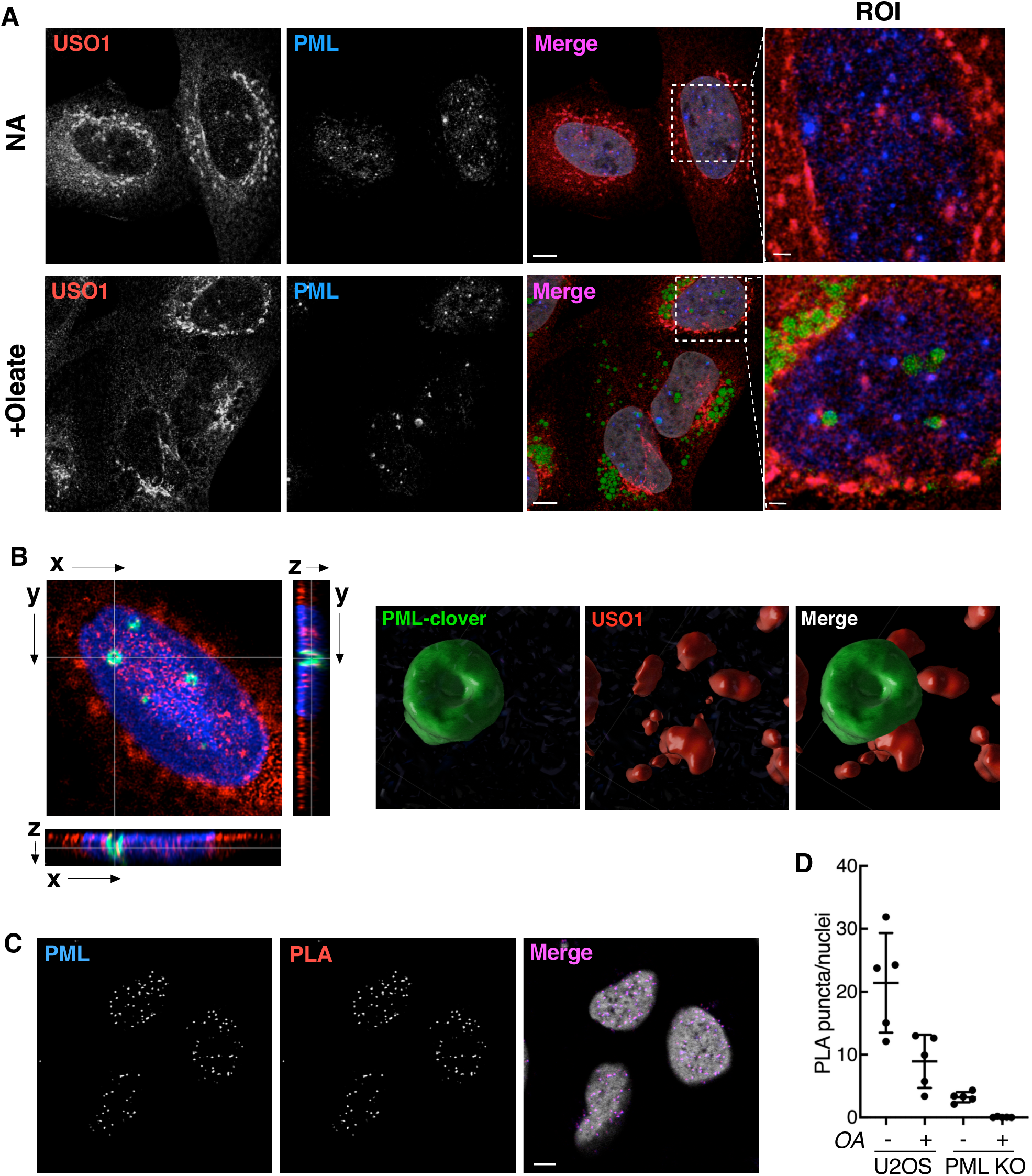
Detection of USO1 interaction with PML NBs and LAPS using immunofluorescence microscopy and proximity ligation assays. A, U2OS cells cultured in the absence or presence of 500 μM oleate for 24 h were immunostained with USO1 and PML primary antibodies followed by AlexaFluor-594 and −647 secondary antibodies, respectively. LDs were visualized with BODIPY493/503 and the nuclei with Hoechst (grey) (bar, 5 μm). The ROI has the Hoechst channel removed (bar, 1 μm). B, xyz projections of confocal sections of a U2OS Clover-PML knock-in cell immunostained for USO1. The nuclei were visualized with Hoechst (blue) and LDs with LipidTOX neutral red. To the left is a 3D reconstruction of the LAPS at the x-y intersection. C, PLA in U2OS cells incubated with PML monoclonal and USO1 polyclonal antibodies. PML was detected with a AlexaFluor647 secondary antibody. D, quantification of PLA nuclear puncta for USO1 and PML in U2OS and PML KO cells incubated in the presence or absence of oleate for 24 h. Results are the means and SD of a representative experiment (5 fields of cells).

Lastly, we investigated whether the expression or nuclear localization of Sec23B, Sec24A or USO1 was affect by PML KO in U2OS cells. Immunoblot analysis showed that expression of all three proteins was unaffected by PML KO or treatment with oleate (Fig. S2A). As well, the nuclear localization of Sec23B and Sec24A in U2OS and PML KO cells was similar (Fig. S2B). However, immunofluorescence localization of USO1 in the nucleus of PML KO cells was reduced compared to controls but cytoplasmic Golgi/vesicular staining was unaffected (Fig. 8A and B). This result is shown more clearly by the quantification of nuclear localized USO1 from images (for example Fig. 8B) and expression as a fraction of total nuclear area (Fig. 8C). It was apparent that the localization of USO1 to the nucleus of U2OS cells was not affected by oleate treatment. However, PML KO cells had a significant reduction in nuclear USO1 whether cultured with or without oleate. Thus, the interaction of USO1 with PML structures appears to be necessary for its retention in the nucleus.

**Figure 8.**
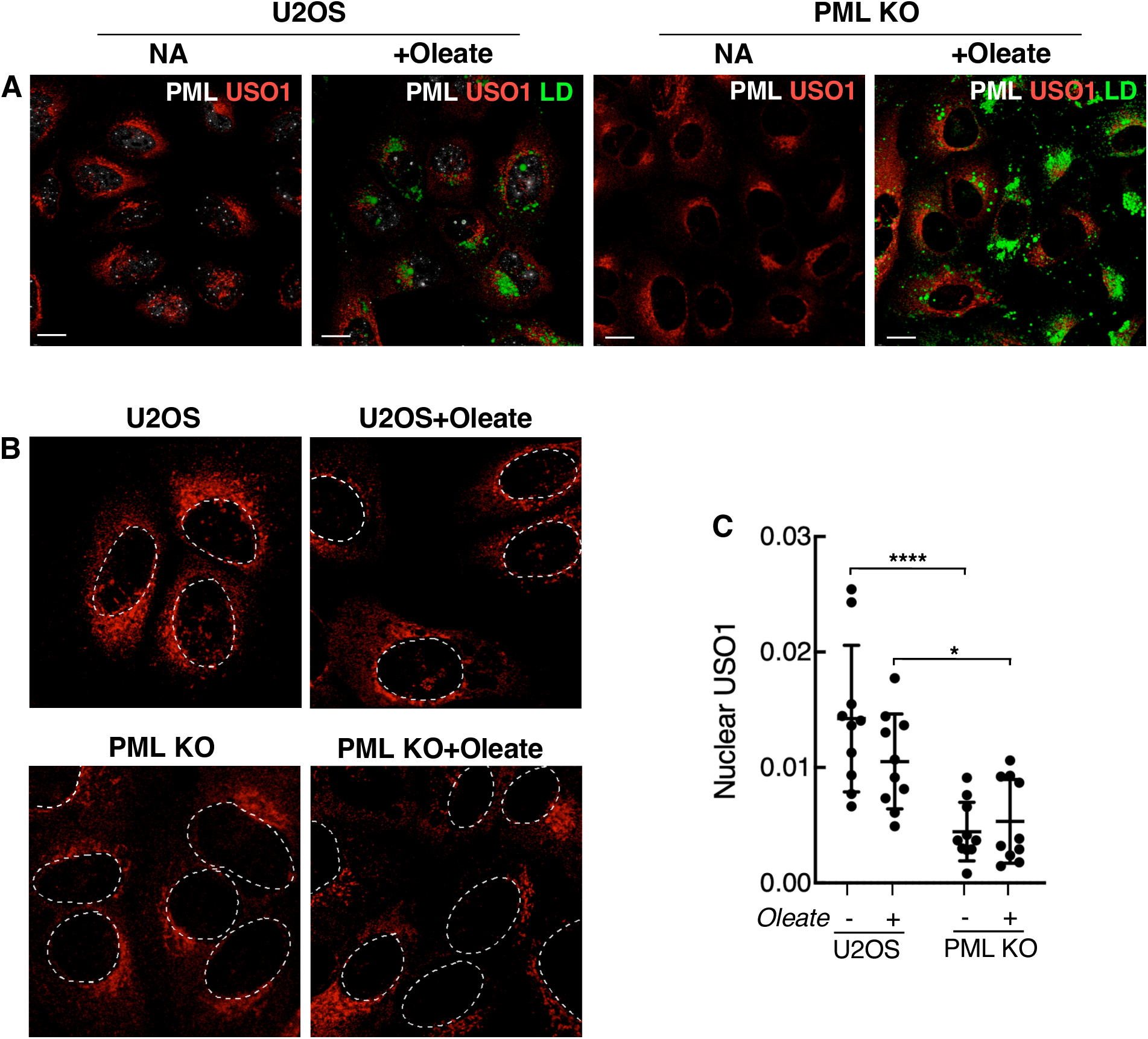
PML knockout U2OS cells have reduced expression of nuclear-localized USO1. A, U2OS and PML KO cells were cultured with no addition (NA) or oleate (500 μM) for 24 h prior to immunostaining with USO1 and PML primary antibodies followed by AlexaFluor-594 and - 647 secondary antibodies, respectively. LDs were visualized with BODIPY493/503 (bar 5 μm). B, cells treated as described above were immunostained for USO1 (nuclei are outlined). C, quantitation of nuclear-localized USO1 expressed as a fraction of total nuclear area. Results are the mean and SD of 10 fields of cells from two independent experiments (\**p*<0.05, \*\*\*\**p*<0.0001).

## DISCUSSION

In response to cellular stressors, PML NBs reorganize into non-canonical structures whose functions are poorly understood due in part to a lack information on their repertoire of interacting proteins. LAPS are one of these alternate PML subdomains formed when PML and PML-client proteins are recruited to the surface of nLDs that form in response to cellular stress induced by excess fatty acids. LAPS are relatively depleted of several PML NB client proteins such as SP100 and DAXX but are enriched in enzymes involved in lipid synthesis and storage, indicating a dramatic shift in PML function to cope with excess fatty acids (Lee et al., 2020). Our understanding of LAPS function and relationship to PML NBs is limited in scope and biased toward proteins with known or specific biochemical functions. To circumvent this issue, we conducted an unbiased APEX2 proximity analysis of PMLI and PMLII interactors in cells treated with oleate to promote LAPS formation compared to untreated cells harbouring only PML NBs. We chose APEX2 in part due to its smaller mass compared to biotin ligases, rapid catalysis for better temporal resolution of bait-protein interactions, and synthesis of a short-lived biotin phenol radical that reacts with electron-rich amino acid residues (i.e. Tyr, Trp, Cys, His) (Lam et al., 2015; Trinkle-Mulcahy, 2019). Ours is the first reported APEX2 proximity labelling study for PML and therefore yielded a unique interactome that is relatively devoid of known PML NB- and LAPS-associated proteins. Indeed, cross referencing the APEX2-PML interactome against the BioGrid data base (Oughtred et al., 2019) for PML revealed common partners from only two studies; a split SUMO-TurboID screen using PML as a substrate (LPP, PPP1R13L, STAM, SEC24B, TNKS1BP1, IGBP1 and ERCC6L) (Barroso-Gomila et al., 2021) and an affinity capture-western study (CSNK1D) (Alsheich-Bartok et al., 2008). Thus, the novel set of APEX2-PML interactors identified in our study reflects the choice of experimental system (ie. cells and treatments) as well as features of the APEX2 probe.

Initially we expected the enrichment of PML-interacting proteins in oleate-treated cells would define a subset of proteins that selectively associate with LAPS; however, our assessment of the interactors that fell into this group suggested otherwise. For example, Sec24B, USO1 and MK2 were enriched by oleate treatment but immunofluorescence microscopy and PLA indicted they associated primarily with PML NBs. An explanation for this discrepancy could lie with the substrate biotin phenol, which is more hydrophobic than biotin and could preferentially accumulate in the hydrophobic lipid environment of LAPS leading to enhanced labeling of proteins in those structures relative to PML NBs. Thus, biotinylation under oleate-treated conditions is not an exhaustive method for the identification of proteins with preferred association with PML NBs or LAPS during lipid stress. As such orthogonal studies, including analysis of PML-client protein and lipid enzyme interactomes using affinity purification methods and different BirA and APEX2-mediated proximity proteomic methods will be required to enable a more thorough characterization of the lipid stress-induced proteome of PML-containing nuclear subdomains.

The interaction of MK2 with PMLI and PMLII in oleate-treated cells attracted interest based on the recent finding that MK2 phosphorylation by p38, and subsequent suppression of necroptosis by phosphorylation of RIPK1 S321 phosphorylation, is negatively regulated by sequestration of both kinases in PML NBs (Chen et al., 2021). Immunofluorescence microscopy showed that the MK2-II isoform was weakly associated with cLDs and LAPS in U2OS cells but attempts to confirm interactions by PLA or other methods were unsuccessful. Oleate-induction of LAPS and nLDs caused a minor but significant reduction in the phosphorylation and activation of p38 in PML KO cells that did not translate into a reduced phosphorylation of its downstream targets MK2 and HSP27. These data suggest that p38 phosphorylation is partially PML-dependent and subject to inhibition by oleate if PML is absent. Thus, the importance of the p38-MK2 pathway in lipid stress remains unclear and may require alternative cell models or modalities of p38 activation to elucidate the significance of this pathway in the context of LAP formation and function.

A novel finding in the current study was the identification of two nodes of PML-interacting proteins involved in cytoplasmic COPII and ESCRT vesicle trafficking. ESCRT complexes in endosomes recruit ubiquitinated proteins into multivesicular bodies that are degraded upon fusion with primary lysosomes (Futter et al., 1996). ESCRT-1 proteins TSG101, ESCRT-0 proteins STAM, STAM2 and HGS, and the ESCRT-associated phosphatase PTPN23 and NEDD4 ligase were identified in both PMLI and PMLII interaction sets. STAM was independently identified in a recent split SUMO-TurboID screen (Barroso-Gomila et al., 2021). In conjunction with VPS13D, TSG101 remodels the phospholipid monolayer of cLDs to assist in the removal of fatty acids to the mitochondria for oxidation during glucose starvation (Wang et al., 2021). TSG101 has been localized to the nucleus (Sun et al., 1999) where it could have a related function on nLDs or LAPS.

Non-canonical functions of COPII vesicle transport proteins Sec23B, Sec24A, Sec24B, Sec23IP, TRAPPC3 and USO1 are also inferred from the APEX2-PML interactome. Immunofluorescence microscopy and PLA confirmed the interaction of the COPII coat proteins Sec23B and Sec24A, as well as the Golgi tethering factor USO1, with PML NBs. Similar to the canonical PML NB client protein DAXX, all three COPII transport proteins had diminished PLA puncta in oleate-treated cells suggesting they dissociate from PML NBs during or after remodelling into LAPS. In contrast to CCTα, which associates strongly with LAPS, there were no prominent PLA signals for Sec23B, Sec24A or USO1 surrounding LAPS. However, USO1 was identified on LAPS by 3D reconstructions of confocal sections, and its nuclear localization was dependent on PML expression. Overall, these COPII proteins behave like the PML client proteins SUMO, DAXX and SP-100 (Lee et al., 2020), which dissociate from PML NBs during the transition to LAPS. A caveat of this interpretation is that immunofluorescence imaging and PLA using Sec23B and Sec24A antibodies required methanol fixation, which could disrupt PML NBs and LAPS. As a result, the PLA signal for Sec23B and Sec24A was reduced (Fig. 6E and G; 2-3 puncta per cell) compared to PLA using CCTα or DAXX antibodies (Fig. 6A and C; 7-10 puncta per cell). Disruption by methanol would cause underestimation of Sec23B and Sec24A interactions with PML NBs and LAPS.

Studies on Sec23B and USO1 suggest some intriguing connections to nuclear PML NB and/or LAPS function. COPI and COPII components, including Sec23 proteins, facilitate the delivery of proteins to the surface of cLDs by forming bridges or fusion events at the ER or ER-Golgi intermediate compartment (Soni et al., 2009; Wilfling et al., 2014). The presence of Sec23B and other COPII components in the nucleus suggests a parallel activity on LAPS. Furthermore, Sec23B is specifically localized to the nucleoli where it is involved in the ER-stress response to proteosome inhibition and transcription of ribosomal protein genes (Yehia et al., 2018; Yehia et al., 2021). This observation raises the possibility that Sec23B and possibly other COPII proteins could mediate an ER stress response induced by fatty acid overload. Nuclear-localized USO1 has been identified as a potential RNA binding protein that is dysregulated in B-cell acute lymphoblastic leukemia and multiple myeloma (Jaiswal et al., 2021; Castello et al., 2016; Jin et al., 2016). Intriguingly, caspase cleavage of USO1 generates a C-terminal fragment that is SUMOylated and translocated to the nucleus where it enhances the apoptotic response in cells (Mukherjee et al., 2009). Our results showing reduced USO1 in the nucleus in PML KO cells implies that SUMOylation of full-length USO1 could mediate its interaction with PML NBs and/or LAPS as part of yet to be defined stress response pathway.

Our BioID study is the first to utilize APEX2-PML constructs to interrogate the interactome of PML NBs and LAPS, which revealed non-conical nuclear functions for cytoplasmic vesicle transport proteins. Validation of three of these COPII-associated proteins interactions indicated a preferred interaction with PML NBs suggestive of novel moon-lighting functions that are linked to reorganization of PML during fatty acid-induced stress.

## MATERIALS AND METHODS

### Cell culture and transfection

U2OS, U2OS PML knockout (Attwood et al., 2020) and U2OS Clover-PML knock-in cells (Pinder et al., 2015) were cultured in DMEM containing 10% FBS at 37°C in a 5% CO2 atmosphere. To induce the formation of TAG-rich lipid droplets, cells were incubated with a oleate/BSA complex (6.6/1, mol:mol) at the concentrations indicated in figure legends (Goldstein, 1983). Cells were transfected with plasmids using Lipofectamine 2000 at a DNA (μg): Lipofectamine (μl) ratio of 1:3 for 48 h prior to the start of experiments. Cells were treated with stock solutions of SB-203580, PF-3644022 and anisomycin dissolved in DMSO.

### Construction of APEX2 expression vectors

pAPEX2-HA-PMLI and -PMLII were constructed by PCR amplification of the APEX2-HA fragment from pUC57-APEX2-HA (Sultana et al., 2020) using forward and reverse primers containing Nhe1 and Bsrg1 restriction sites, respectively. The APEX2-HA fragment was ligated into pEGFP-C1-PMLI and pEGFP-C1-PMLII (Pinder et al., 2015) that were digested with the same enzymes to remove the GFP cassette. pAPEX2-HA-NLS was constructed by PCR amplification of APEX2-HA fragment from pUC57-APEX2-HA using forward and reverse primers containing Kpn1 and Bsrg1 restriction sites, respectively. The APEX2-HA fragment was cloned into pEGFP-NLS-N1 (3 NLS cassettes inserted at the Bsrg1 site) that was digested with Kpn1 and Bsrg1 to remove the GFP cassette.

### Biotin-phenol labelling of cells expressing APEX2 fusion proteins

U2OS cells transiently transfected with pAPEX2-HA-PML-I, pAPEX2-HA-PML-II, or pAPEX2-HA-NLS were treated with or without 500 μM oleate for 24 h. Cells were then incubated with 0.5 mM biotin-phenol (in DMSO) for 30 min at 37°C in a 5% CO2 incubator, hydrogen peroxide (1 mM) was added for 1 min at room temperature, and the reaction terminated by the addition of 0.5 ml of quench solution (10 mM Trolox, 20 mM ascorbate in PBS, pH 7) (Rhee et al., 2013). Cells were rinsed twice with 0.5 ml of quench solution and lysed for SDS-PAGE, fixed for immunofluorescence or isolated on NeutrAvidin-agarose (Thermo-Fisher) for mass spectrometry of biotinylated peptides (see below).

### Isolation of biotinylated proteins on NeutrAvidin-agarose

After proximity biotin labelling as described above, cells were scraped in PBS, collected by centrifugation at 3,000*xg* for 5 min and the cell pellet was dissolved for 5 min on ice in 1 ml of 50 mM Tris (pH 7.4), 5 mM Trolox, 0.5% (w/v) sodium deoxycholate, 150 mM NaCl, 0.1% (w/v) SDS, 1% (v/v) Triton X-100, 1 mM phenylmethanesulfonylfluoride, 10 mM NaN3, 10 mM sodium ascorbate, and protease inhibitor cocktail (James et al., 2019). Cell lysates were centrifuged for 10 min at 13,000*xg* at 4°C and the supernatant collected and incubated with NeutrAvidin-agarose for 12 h at 4°C on an orbital rotor. The beads were collected by brief centrifugation, washed once with buffer 1 (50 mM HEPES, pH 7.4, 0.1% (w/v) sodium deoxycholate, 1% (v/v) Triton X-100, 500 mM NaCl, and 1 mM EDTA), once with buffer 2 (50 mM Tris, pH 8.0, 250 mM LiCl, 0.5% (v/v) Nonidet P40, 0.5% (w/v) sodium deoxycholate, and 1 mM EDTA), and twice with buffer 3 (50 mM Tris, pH 7.4, 50 mM NaCl). During each step the beads were incubated for 5 min at 4°C on an orbital rotor. The proteins were eluted from the agarose beads by incubation with SDS lysis buffer supplemented with 5 mM desthiobitoin for 5 min at 95°C. Protein samples were then resolved by SDS-PAGE and immunoblotted to detect biotinylated proteins (see below).

### Identification of biotinylated proteins by quantitative mass spectrometry

Sample preparation and analysis was conducted as previously described (Del Olmo et al., 2019). Proteins bound to NeutrAvidin-agarose were subject to 5 washes with 20 mM NH4CHO3, reduced with 10 mM DTT for 30 min and alkylated with 15 mM iodoacetamide for 1 h. On-bead digestion with trypsin (1 μg overnight) was stopped with 1% formic acid. Eluted peptides were collected, and the beads were treated with 1% formic acid and 60% acetonitrile to release residual peptides. Pooled eluates were dried, resuspended in 0.1% formic acid and 2 μg was injected into an HPLC (nanoElute, Bruker Daltonics) equipped with a trap column (Acclaim PepMap100 C18, 0.3 mm id x 5 mm Dionex Corporation) and analytical C18 column (1.9 µm beads, 75 µm x 25 cm, PepSep). Peptides were separated over 4 h using a linear gradient of 5-37% acetonitrile in 0.1% TFA at a flow rate of 500 nL/min while being injected into a TimsTOF Pro ion mobility mass spectrometer equipped with a Captive Spray nano electrospray source (Bruker Daltonics). Data was acquired using data-dependent auto-MS/MS with a 100-1700 m/z mass range, with parallel accumulation-serial fragmentation (PASEF) enabled with a number of PASEF scans set at 10 (1.27 seconds duty cycle) and a dynamic exclusion of 0.4-minute, m/z dependent isolation window and collision energy of 42.0 eV. The target intensity was set to 20,000, with an intensity threshold of 2,500.

The raw files were analyzed using MaxQuant version 1.6.17.0 software (20) and the Uniprot human proteome database (21/03/2020, 75,776 entries). The settings used for the MaxQuant analysis (with TIMS-DDA type in group-specific parameters) were: 2 miscleavages were allowed; fixed modification was carbamidomethylation on cysteine; enzymes were trypsin (K/R not before P); variable modifications included in the analysis were methionine oxidation, protein N-terminal acetylation and protein carbamylation (K, N-terminal). A mass tolerance of 10 ppm was used for precursor ions and a tolerance of 20 ppm was used for fragment ions. Identification values “PSM FDR”, “Protein FDR” and “Site decoy fraction” were set to 0.05. Minimum peptide count was set to 1. Label-Free-Quantification (LFQ) was also selected with a minimal ratio count of 2. Both the “Second peptides” and “Match between runs” options were also allowed. Following the analysis, the results were sorted according to several parameters. Proteins positive for at least either one of the “Reverse”, “Only.identified.by.site” or “Potential.contaminant” categories were eliminated, as well as proteins identified from a single peptide.

The average peptide count data was filtered using the CRAPome repository (Mellacheruvu et al., 2013) with a threshold score of 0.2. to remove common mass spectrometry contaminants. The residual proteins from the APEX2-PMLI and -PMLII data sets were then compared against the APEX2-NLS protein data sets using the default settings of the program SAINTexpress using a score threshold of 0.2 (Teo et al., 2014). Dot plots of the resulting proteins were made using ProHits-viz with a Bayesian false discovery rate threshold of 0.05 (Knight et al., 2017) and interaction networks analyzed using STRING v11 (Szklarczyk et al., 2019).

### Immunoblotting

Cells were rinsed and scraped in cold PBS, collected by centrifugation at 10,000*xg* for 10 mins at 4°C, and the cell pellet lysed with SDS lysis buffer (62.5 mM Tris-HCl, pH 6.8, 10% glycerol, 2% SDS, 0.05% bromophenol blue and 5% β-mercaptoethanol) on ice for 10 min. Cell lysates were sonicated for 10 sec, denatured at 90°C for 3 min, resolved by SDS-PAGE and transferred to nitrocellulose filters. After incubation in LI-COR Odyssey blocking solution diluted 4:1 (v/v) in TBS-Tween (20 mM Tris HCl, pH 7.4, 150 mM NaCl, 0.1% Tween 20) for 1 h at 20°C, the filters were incubated with the following primary antibodies; HA (mouse monoclonal 2367S, Cell Signalling), PML (rabbit polyclonal A301-167A, Bethyl), MK2 (rabbit polyclonal PA5-17729, Invitrogen), p38 (rabbit polyclonal 9212S, Cell Signalling), HSP27 (rabbit polyclonal 2402T, Cell Signalling), p38-pT180/Y182 (rabbit polyclonal 9216S, Cell Signalling), MK2-pT334 (rabbit polyclonal 3007T, Cell Signalling) or HSP27-pS82 (rabbit polyclonal 9709T, Cell Signalling). After rinsing in TBS-Tween, the filters were incubated with goat anti-mouse and goat anti-rabbit secondary antibodies conjugated to LI-COR680 or 800 IR dyes, or streptavidin-conjugated LICOR680 IR for 1 h at 20°C. Nitrocellulose membranes were imaged and fluorescence quantified using a LI-COR Odyssey scanner and associated software (v3.0).

### Immunofluorescence confocal microscopy

Cells cultured on glass coverslips (0.13-0.16 mm) were fixed with paraformaldehyde (4%, w/v) in PBS for 15 min at 20°C, permeabilized with Triton X-100 (0.1%, v/v) in PBS for 10 min at 4°C and blocked with BSA (1%, w/v) in PBS (PBS/BSA) for 1 h at 20°C. For immunostaining with Sec23B and Sec24A antibodies, cells were fixed in 100% methanol at −20°C for 2 minutes and blocked with PBS/BSA. Coverslips were incubated with the following primary antibodies diluted in PBS/BSA overnight at 4°C; FLAG (mouse monoclonal F1804, Sigma Aldrich), Sec24A (rabbit polyclonal HPA056825, Sigma Aldrich); Sec23B (rabbit polyclonal HPA008216, Sigma Aldrich); USO1 (rabbit polyclonal 13509-1-AP, Proteintech) PML (mouse monoclonal sc-377390, Santa Cruz) as well as those listed in the previous section. After washing in PBS/BSA, coverslips were incubated with goat anti-rabbit and goat anti-mouse secondary antibodies conjugated to AlexaFluor-488, −594 or −647 or streptavidin-AlexaFluor-488. LDs were detected with BODIPY493/503, LipidTOX neutral red or LipidTOX deep red diluted in PBS. Nuclei were visualized with Hoescht. Coverslips were mounted in Mowiol on glass slides and imaged using a Leica SP8 LIGHTNING confocal microscope with a 63x oil immersion HC Plan APOCHROMAT objective, 405 nm, 488 nm, 552 nm and 638 nm lasers, and Leica LAS X software. For 3D renderings, three-channel confocal Z-stacks of USO1 Clover-PML cells immunostained for USO1, LipidTOX neutral red and Hoechst were captured using a Leica SP8 LIGHTNING. Z-stacks were rendered using Imaris software (9.7.0) and regions of interest were cropped as previously described (Dorighello et al., 2023).

To quantify nuclear USO1 area, images were converted to 8-bit and a tailored threshold applied for the Hoechst channel. Total nuclear area using the Hoechst channel was quantified using the Measure function. To quantify USO1 nuclear area, a nuclear mask was applied to the USO1 channel, nuclear USO1 was converted to 8-bit, a uniform threshold applied, and the area was quantified using the Measure function. Nuclear USO1 was expressed as a fraction of total nuclear area.

### Proximity ligation assay

After treatments, U2OS and PML KO cells seeded on coverslips (0.13-0.16 mm) were fixed and permeabilized as described in the previous section. Cells were incubated with primary mouse/rabbit antibody pairs overnight at 4°C in PBS/BSA: CCTα rabbit polyclonal (Gehrig et al., 2008), DAXX (rabbit polyclonal D7810, Sigma Aldrich) as well as those listed in the previous sections. PLA were performed according to the manufacturer’s instructions (DuoLink Red kit, Sigma-Aldrich). After removing primary antibodies, PLA anti-rabbit PLUS and anti-mouse MINUS secondary antibodies (diluted in 40 μl PBS/BSA) were spotted on parafilm in a humidified chamber, and coverslips were inverted on the droplets for 1 h at 37°C. After removing the secondary antibodies, ligation and amplification reactions using fluorophore-labelled oligonucleotides (λEx593 nm/λEx 622) were performed in 40 μl reaction mixtures at 37°C for 30 and 100 min, respectively. To detect the primary antigens, coverslips were occasionally incubated with anti-rabbit or anti-mouse secondary antibodies conjugated to AlexaFluor-488 or −647. Coverslips were dried before being mounted on glass slides using In Situ mounting medium containing Hoechst and sealed with nail polish. To quantify PLA puncta, Image J (v1.53) was used to convert each PLA and PML images to 8-bit and a uniform threshold applied to each channel. The number of PML positive puncta was quantified from these two channels using the AND Image Calculator and Analyze Particles function with the default settings. For the CCTα PLA quantification, the Analyze Particles function was used on the PLA channel alone. The nuclei were quantified from the Hoechst channel.

## Abbreviations

BioID: proximity-dependent biotin identification
CCT: CTP:phosphocholine cytidylyltransferase
cLD: cytoplasmic lipid droplet
ER: endoplasmic reticulum
ESCRT: Endosomal sorting complexes required for transport
LAPS: Lipid-Associated PML structure
KO: knockout
nLD: nuclear lipid droplets
MK2: mitogen activated protein kinase-activated protein kinase
PLA: proximity ligation assays
PC: phosphatidylcholine
PLA: proximity ligation assay
PML: promyelocytic leukemia
TAG: triacylglycerol
RBBC: RING-B-box coiled-coil
SIM: SUMO-SUMO interaction motifs.

## ACKNOWLDGEMENTS

We thank Robert Douglas for technical assistance with tissue culture.

